# Remote Cortical Network for Frontal Cognitive Symptoms Derived from Tau Accumulation in Progressive Supranuclear Palsy

**DOI:** 10.1101/2025.02.05.636578

**Authors:** Yuki Hori, Hironobu Endo, Kenji Tagai, Yuko Kataoka, Ryoji Goto, Shin Kurose, Yuki Momota, Naomi Kokubo, Chie Seki, Sho Moriguchi, Hitoshi Shimada, Hitoshi Shinotoh, Takahiko Tokuda, Keisuke Takahata, Takafumi Minamimoto, Makoto Higuchi, Toshiyuki Hirabayashi

## Abstract

Progressive Supranuclear Palsy (PSP) is a neurodegenerative disorder characterized by impaired movement and frontal cognitive dysfunctions. While motor symptoms have been linked with subcortical tau deposits, network mechanisms underlying the frontal cognitive symptoms remain unclear because pathological tau accumulation is quite limited within the frontal cortex and heterogeneously distributed primarily in the subcortical regions. Here, we combined tau-PET using a high-contrast probe we have recently developed with normative connectome to resolve this issue. Tau-deposition sites were connected to a common cortical network that could not be identified through similar analyses based on atrophy detected by structural MRI. This network was predominantly overlapped with canonical action-mode and frontoparietal networks, which jointly support adaptive and goal-directed behavior. Critically, while the degree of subcortical primary tau deposition correlated with motor symptoms, only the degree of connectivity from tau-deposition sites to the derived cortical network explained the frontal cognitive deficits of individual patients. These findings suggest a novel mechanism that frontal cognitive impairments, but not motor deficits, in PSP are derived from remote effects of tau deposits, independent of atrophy, through the convergent connectivity to the identified common cortical network.

## INTRODUCTION

Progressive supranuclear palsy (PSP) is a neurodegenerative disorder primarily characterized by movement impairments such as oculomotor deficits, postural instability, and akinesia (Hauw et al., 1994; Hoglinger et al., 2017; Steele et al., 1964). In addition to these motor-related symptoms, PSP is frequently associated with cognitive deficits, especially those related to frontal cortical functions (Donker Kaat et al., 2007; Dubois et al., 2000). Pathologically, PSP is known as a four-repeat tauopathy, where the aggregation of hyperphosphorylated tau proteins is implicated as a major cause of clinical symptoms, accompanied by neuroinflammation and atrophy (Ballatore et al., 2007; Maruyama et al., 2013). Tau lesions, neuronal loss, and gliosis have been found primarily in subcortical regions and in a part of cortical regions, including the basal ganglia, midbrain, motor-related cortical areas, and cerebellum (Verny et al., 1996), and damage to these structures has been associated with motor dysfunction and general disease severity (Alster et al., 2019; Hauw et al., 1994; Hauw et al., 1990; Hoglinger et al., 2017). Subcortical atrophy— especially in the midbrain and cerebellum—is another notable pathological hallmark of PSP, and can be used to distinguish patients with PSP from those with Parkinson’s disease or age-matched healthy controls (Chen et al., 2024; Price et al., 2004). However, the mechanisms underlying the frontal cognitive decline in PSP remain elusive, as atrophy or tau deposition within the frontal cortex itself is only rarely observed beyond the primary and supplementary motor areas (Boxer et al., 2006; Cope et al., 2018; Josephs et al., 2008; Price et al., 2004).

Neurodegenerative diseases, including PSP, are increasingly being conceptualized as network-based disorders (Seeley et al., 2009). The aggregation of pathological proteins in specific brain regions disrupts synaptic function, with damage propagating across interconnected networks (Graveland et al., 1985; Seeley et al., 2006), and accelerating clinical symptoms (Selkoe, 2002). In PSP, reduced functional connectivity (FC) between the atrophied midbrain region and other areas including the frontal cortex has been reported (Gardner et al., 2013). In addition, hypoperfusion (Gurd and Hodges, 1997; Roemer et al., 2024) and hypometabolism (Salmon et al., 1997; Whitwell et al., 2017) in the frontal cortex have been observed even in the absence of the corresponding local pathological changes, suggesting that the frontal cognitive dysfunctions in PSP might result from the remote effects of subcortical pathology.

Using normative connectome data, lesion network mapping (LNM) is a powerful approach for identifying brain regions commonly connected from heterogeneous lesion sites that are associated with shared symptoms across patients with stroke. This approach is based on the principle that in addition to direct local impairments, a lesion in a given brain region also indirectly impairs functions of remote regions connected from the lesion site. LNM has uncovered hidden core regions associated with diverse symptoms across neurological conditions, including visual hallucinations (Boes et al., 2015), delusional misidentifications (Darby et al., 2017), amnesia (Ferguson et al., 2019), loss of consciousness (Fischer et al., 2016), criminal behavior (Darby et al., 2018), and abnormal movements such as freezing of gait and hemichorea-hemiballismus (Laganiere et al., 2016). Importantly, core regions identified through LNM have been shown to align with previously identified neuromodulatory targets for symptom relief (Joutsa et al., 2022; Padmanabhan et al., 2019), further validating its utility. Recently, LNM has been successfully extended to tau deposition- or atrophy-based analysis in Alzheimer’s disease (AD) (Luan et al., 2024; Tetreault et al., 2020; Zhou et al., 2025), demonstrating its potential for identifying common brain locations functionally connected from regions of pathology in spite of across-patient variability in their spatial patterns in a given neurodegenerative disease. However, tau/atrophy-network mapping has not been applied to non-AD tauopathies, and critical comparisons between the influences of local tau-deposits/atrophy and their remote network effects on a specific symptom, and between the effects of tau-deposits and atrophy on the network mapping have not been conducted yet for any neurodegenerative diseases.

In the current study, we hypothesized that a common network extended from tau deposition is responsible for the frontal cognitive dysfunctions observed frequently in PSP. Specifically, we first examined whether such common network can be identified for PSP, and further investigated whether motor and frontal cognitive impairments in PSP could be attributed to local tau deposition or its remote effects exerted on the identified common extended network. To achieve this, we used florzolotau (18F) (also known as PM-PBB3 or APN-1607), a novel PET probe that we recently developed for detecting tau deposits (Tagai et al., 2021). This probe provides high metabolic stability and remarkable contrast at the single-patient level not only for AD, but also for non-AD tauopathies including PSP, which has been difficult with previous probes (Tezuka et al., 2021). By applying network mapping to PET-detected tau deposits in individual patients, we identified common brain regions that are remote but functionally connected from heterogeneous locations of pathological tau aggregates, and examined their relationships with clinical symptoms (Fig. 1). Our results indicate that a network mechanism underlies the frontal cognitive impairments in PSP and highlight the distinct contribution of remote effects mediated by tau deposits.

**Figure 1.**
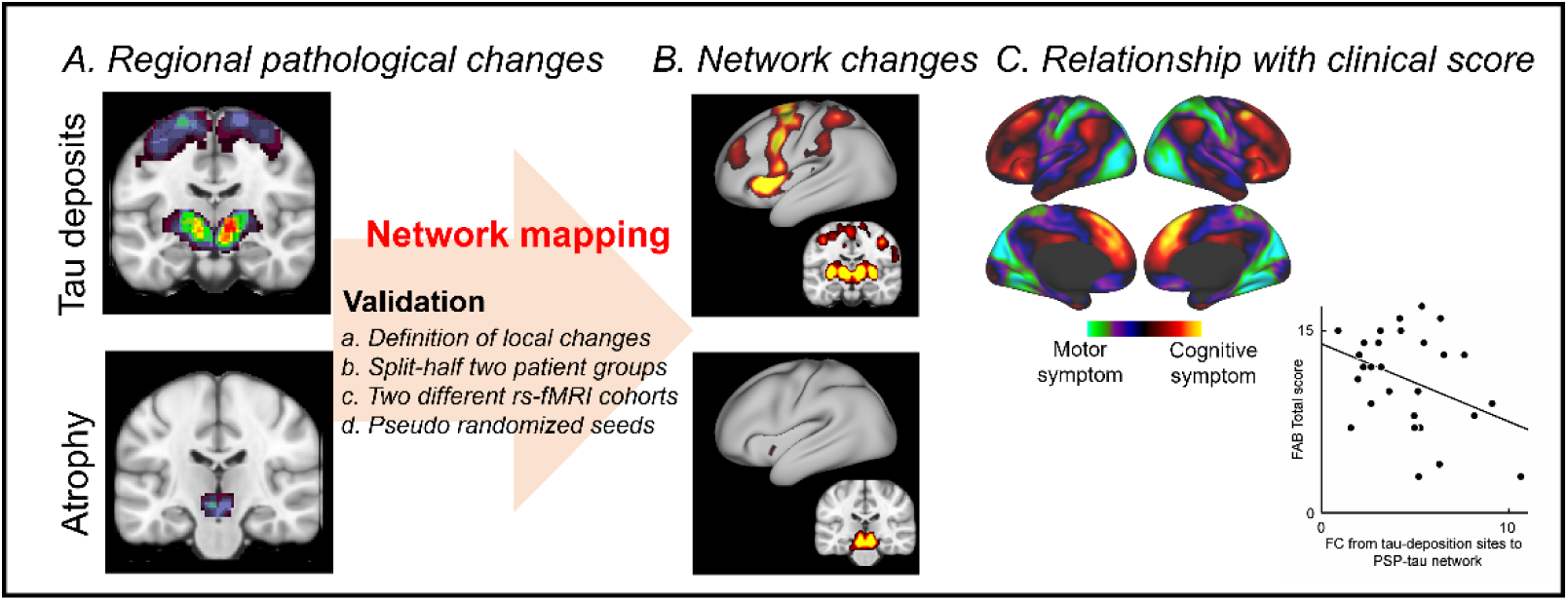
Overview of the study. Regional and network impairments responsible for the frontal cognitive dysfunctions were examined in PSP patients. **(A)** Regional impairments caused by tau deposits and atrophy were identified via ^18^F-florzolotau PET and MRI, respectively. **(B)** Common network impairments across the patients were identified using a resting-state functional MRI (rs-fMRI) database of healthy participants with tau-deposition/atrophy sites in individual patients as seed regions. **(C)** Finally, whether clinical symptoms of individual patients could be best explained by the degree of local pathologies or their network effects.

## RESULTS

### Localization of tau deposits and brain atrophy in individual patients with PSP

We analyzed the data from 37 patients (22 males and 15 females) with PSP-Richardson syndrome, which presents the most typical clinical features of PSP (Endo et al., 2022). The mean patient age and disease duration were 70 ± 1.2 years and 3.4 ± 0.40 years (mean ± SEM, respectively. Disease severity as assessed by the PSP rating scale (PSPRS) was 41 ± 2.9. All patients exhibited tau deposits visualized and quantified with florzolotau PET, without any accumulation of amyloid-beta, as determined by Pittsburgh compound-B (PiB) PET.

To identify the locations of tau deposition and atrophy, we first calculated *t*-score maps of florzolotau PET and MRI-based voxel-based morphometry (VBM) images for each patient. These maps were generated by comparing patient data with the corresponding images of age-matched healthy controls (N = 48) (Fig. 2). Across patients, tau deposits were most frequently observed subcortically in the globus pallidus (GP) (*p* < 0.0003, unpaired *t*-test) and midbrain (*p* < 0.0006), with 62% and 57% overlaps across patients when thresholded at *p* = 0.001, respectively (fig. S1C). In addition, moderate tau accumulation was also detected in motor-related cortical areas in the precentral gyrus (*p* < 0.003, 25% overlap). Meanwhile, brain atrophy was primarily localized to the midbrain (*p* < 0.01, 27% overlap), cerebellum (*p* < 0.03, 27%), and GP (*p* < 0.05, 14%) (fig. S1, B and D). Note that overlap of both tau deposition and atrophy was extremely limited in the frontal cortex, except for some regions in the primary/supplementary motor and premotor areas.

**Fig. 2.**
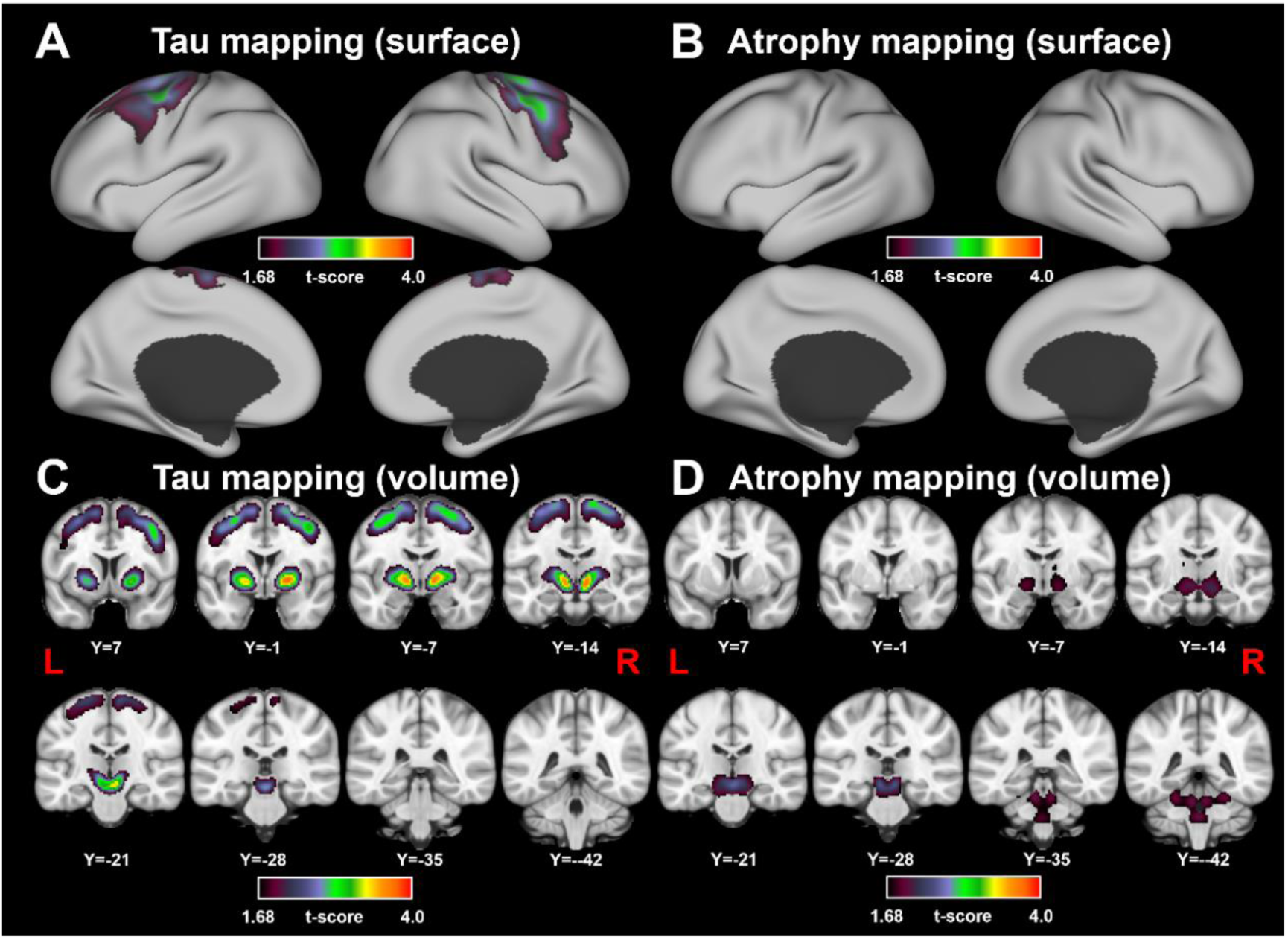
Population maps of tau deposition and atrophy among patients with PSP. **(A-B)** Mean *t*-score maps of tau deposition (A, C) and atrophy (B, D) among PSP patients were presented on the surface (A, B) and volume space (C, D). Tau deposits were primarily observed subcortically in the globus pallidus and midbrain. Moderate accumulation of tau was also detected in motor-related cortical areas in the precentral gyrus. Brain atrophy was observed in the midbrain and cerebellum. For both tau deposit and atrophy, unpaired *t*-test was conducted against age-matched healthy control data.

#### Common extended network derived from tau deposits or brain atrophy across patients with PSP

Next, we sought to determine whether common networks functionally connected from tau-deposit and/or atrophy locations could be observed across the patients. We first examined the brain regions that were functionally connected from the tau deposit locations in each patient using a tau-network mapping approach (see fig. S2). The locations of tau deposit in each patient were identified based on the *t*-map (*p* < 0.001, two-sided, uncorrected). These locations then served as seed regions for calculating the whole-brain FC maps using a publicly available resting-state functional MRI (rs-fMRI) dataset of healthy subjects [Brain Genomics Superstruct Project (GSP) (Yeo et al., 2011)] (N = 100) (Boes et al., 2015). Although tau deposits were restricted to subcortical areas and some portions of motor-related cortex (Fig. 2, A and C), the analysis revealed a common network derived from tau deposits that extended to widespread cortical areas, including the lateral prefrontal cortex (LPFC), dorsal anterior cingulate cortex (dACC), posterior parietal cortex (PPC), and anterior insula (AI) (*p* < 0.05, family-wise error (FWE)-corrected; Fig. 3A and D). In contrast, atrophy network mapping based on the thresholded (*p* < 0.001, uncorrected) atrophy maps from the same patients revealed a common extended network largely restricted to subcortical areas, including the midbrain, pons, and a part of cerebellum (Fig. 3, B and E). These results suggest that the extended, common cortical network observed for the PSP patients can be derived specifically from tau deposits, rather than atrophy.

**Fig. 3.**
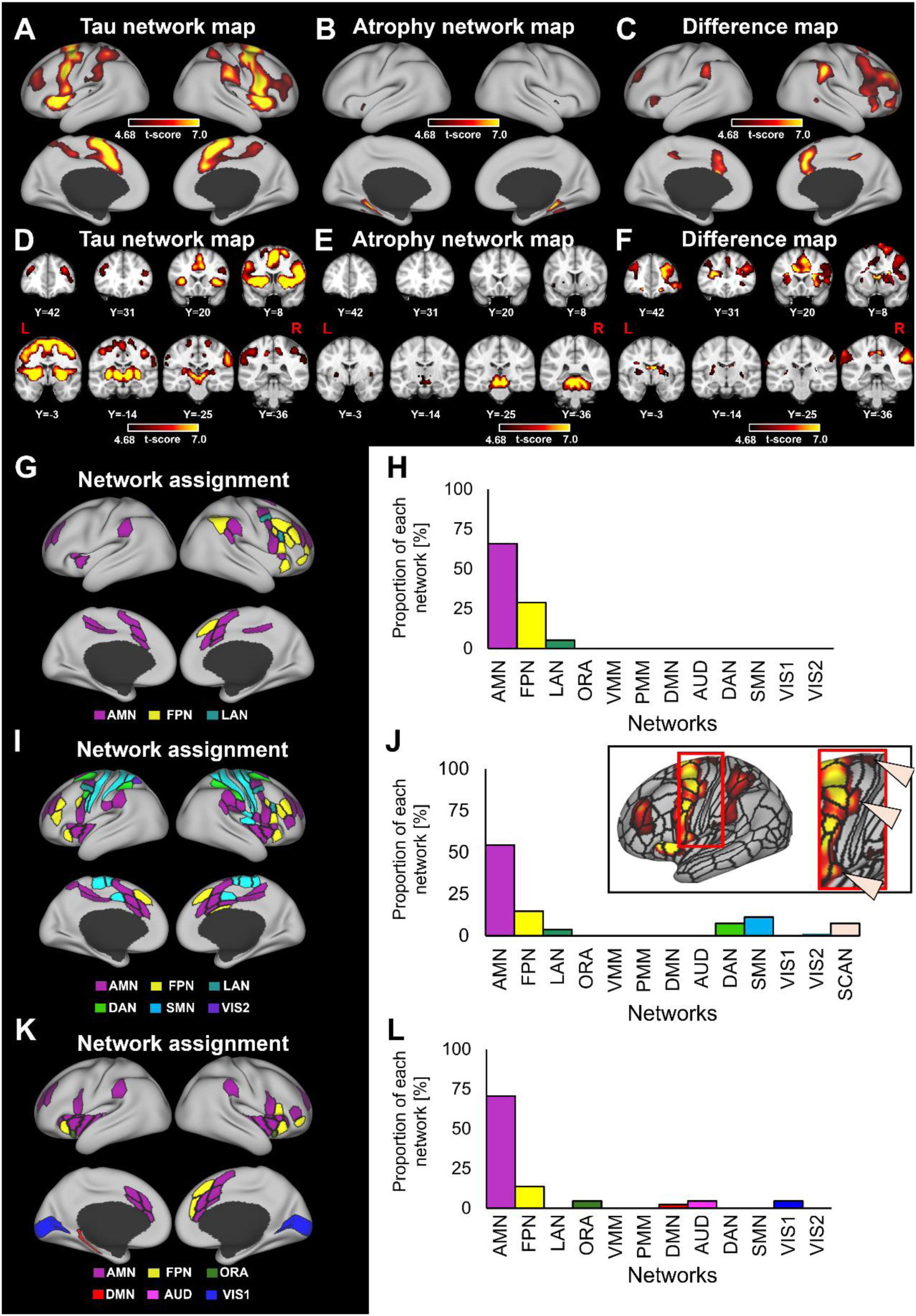
Tau- and atrophy-network maps for patients with PSP. **(A-F)** Average network maps calculated using tau deposit and atrophy VOIs in individual patients as seed regions were presented on the surface (A and B for tau and atrophy, respectively) and volume space (D and E for tau and atrophy, respectively). Difference map was obtained by comparing FC maps seeded from tau deposition sites and atrophy sites (C and F). **(G-L)** Overlap analysis with canonical functional networks. Based on the cortical parcellation in Ji et al. (2019), one of twelve canonical networks was assigned to each region in difference map (G), tau network map before comparison with atrophy-network map (I), and tau-network map calculated from subcortical tau deposits (K). Proportion of each network area to the total area was calculated for difference map (H), tau-network map before comparison with atrophy-network map (J), and tau-network map calculated from subcortical tau deposits (L). In the upper right of panel J, tau-network map focusing on the motor cortex is shown, with three triangles indicating SCAN regions. *AMN: Action-mode network; FPN: Frontoparietal network; LAN: Language network; ORA: Orbito-affective network; VMM: Ventral multimodal network; PMM: Posterior multimodal network; DMN: Default mode network; AUD: Auditory network; DAN: Dorsal attention network; SMN: Somatomotor network; VIS1: Visual network 1; VIS2: Visual network 2; SCAN: Somato-cognitive action network*.

To validate these results, we conducted several control analyses as follows. First, we confirmed that the observed common network was basically conserved even when different thresholds (*p* < 0.005, uncorrected or *p* < 0.05, FWE-corrected) were used to define the seed regions from the tau-accumulation (fig. S3, A-F) and atrophy (fig. S3, G-L) maps, suggesting that the results did not depend on a specific threshold value for seed definition. Second, reproducibility of the common cortical network was assessed by randomly half-splitting the original dataset into two subgroups. The resultant FC maps derived from tau deposits or atrophy were significantly correlated between the two subgroups (fig. S4, A-C and E-G; *r* = 0.87 and 0.73, *p* < 0.001 for tau- and atrophy-network mapping, respectively). We further randomly divided the dataset into two subgroups 100 times. The obtained mean correlation coefficients were significant for both mappings (*r* = 0.92 and 0.88, *p* < 0.001 for tau- and atrophy-network map, respectively), well exceeded the 95th percentile of the correlation coefficients calculated from shuffled pairs of FC values (fig. S4, D and H). The results indicate high and significant reproducibility of the obtained network maps. Third, to rule out the possibility of connectome-specific effects, we calculated the tau deposit-derived network using an alternative publicly available rs-fMRI database [human connectome project (HCP)] (Van Essen et al., 2013). The cortical nodes identified in this analysis were similar to the original ones derived using the GSP database (fig. S5A), and the resultant networks were significantly correlated with each other (Spearman’s Ranked *rho* = 0.30, *p* < 0.001), supporting the robustness of our findings across different rs-fMRI datasets. Lastly, to ensure that the resultant networks were specific to the tau-deposit or atrophy locations and not merely reflected “hub” regions in the brain, we contrasted the tau- and atrophy-network maps against corresponding control maps derived from spatially randomized seeds with the same number of voxels as those of the original tau depositions across the patients. The results were consistent with the original tau- and atrophy-network maps (fig. S6), denying the possibility of reflecting hub regions on the network maps. These results confirmed that the observed common networks were specifically and reliably derived from tau-deposit and atrophy locations in patients with PSP.

Because tau deposits are thought to cause neuronal loss and subsequent atrophy (Ballatore et al., 2007), one possible concern is that the observed tau deposit-derived network might partially reflect the remote effects of atrophy. To isolate the tau-specific extended common network, we further contrasted the tau- and atrophy-derived network maps from the same patients. This analysis revealed predominantly cortical nodes with significantly stronger connectivity from the tau-deposit locations than from the atrophy locations, including the anterior dorsolateral PFC (adlPFC), dorsomedial frontal cortex (dmFC), dACC, premotor cortex (PMC), inferior frontal cortex (IFC), AI, and PPC (*p* < 0.05, FWE-corrected; Fig. 3, C and F). Importantly, no regions showed the opposite pattern (i.e., significantly stronger connectivity from the atrophy locations than from the tau-deposit locations), and most of the above cortical nodes did not exhibit significant tau deposition. We refer to this cortical network derived specifically from tau deposits as the PSP-tau network.

To assess whether the derived PSP-tau network corresponds to specific canonical functional networks in healthy individuals, we mapped the regions in the PSP-tau network onto 12 functional networks defined in Ji et al (Ji et al., 2019) (Fig. 3G). The distribution of regions across the networks were significantly uneven (*p* < 1.0×10^−16^, Chi-squared test), with the majority of regions were located in the action-mode (cingulo-opercular) network (AMN, 66%) (Dosenbach et al., 2025), followed by the frontoparietal (FPN, 29%), and language (LAN, 5.3%) networks. No nodes from any of the other nine networks (orbito-affective, ventral multimodal, posterior multimodal, default mode, auditory, dorsal attention, somatomotor, primary visual, and secondary visual networks) were included in the PSP-tau network (Fig. 3H). These results suggest that the cortical remote dysfunction caused by tau deposition in patients with PSP is predominantly linked to multiple but specific canonical networks, especially AMN and FPN jointly supporting adaptive and goal-directed behavior (Marek and Dosenbach, 2018). Similar pattern was observed for the tau-network map before contrasting with the atrophy-network map (Fig. 3I and J; *p* < 1.0×10^−29^, Chi-squared test; AMN and FPN were the primary and secondly network, respectively). The somato-cognitive action network (SCAN), which is strongly interconnected with the AMN (Gordon et al., 2023), but not defined in (Ji et al., 2019), was also included in the tau-network map, but only before contrasting with atrophy-network map (Fig. 3J, top-right inset). Critically, we also confirmed that a similar network assignment was obtained when seed regions were restricted to subcortical tau deposits across all patients (Fig. 3K and 3L; *p* < 1.0×10^−24^, Chi-squared test; AMN and FPN were the primary and secondly network, respectively), suggesting that the PSP-tau network was derived not only from the cortical tau deposits.

#### FC strength from tau-deposition sites to the PSP-tau network explained across-patient variability of frontal cognitive decline

To assess how the derived PSP-tau network is involved in specific clinical symptoms at the single-patient level, we examined the relationship between clinical scores and the FC strength from tau-deposition sites to the PSP-tau network in the same patients, a proxy of estimated remote effect of tau deposition exerted on the network. In particular, we focused on whether such a remote effect explains frontal cognitive symptoms. The FC values from tau-deposition sites to the PSP-tau network significantly explained the variance in the Frontal Assessment Battery (FAB) score across the patients, a general clinical test for measuring cognitive functions of the frontal cortex (Dubois et al., 2000) (Fig. 4A) (*p* < 0.02). The result was consistent when multiple regression analysis was conducted with the degree of tau deposition in the same network and the local degree of tau deposition in the GP, where both the maximum overlap and the highest *t*-value of tau deposition were observed across the patients (Fig. 4 A), as additional explanatory variables (*p* < 0.02). Conversely, these additional variables themselves did not explain the FAB total score (*p* > 0.1). More closely, FC values from tau-deposition sites to the dmFC node of the PSP-tau network in individual patients, a primary node in the frontal cortex without significant tau deposition (Fig. 3C), significantly explained the FAB total score (*p* < 0.03, FDR-corrected for multiple comparisons) and its subscore Go/NoGo task performance (*p* < 0.05) (Fig. 4B, red), which requires response inhibition and whose decline is well known as a primary symptom of PSP patients (Migliaccio et al., 2020; Zhang et al., 2016). In addition, FC to this node also significantly explained the FAB subscore of conceptualization (*p* < 0.03, FDR-corrected), which has been causally linked with the dPFC dysfunction (Kopp et al., 2013). Meanwhile, the same FC values did not explain more general cognitive abilities indexed by Mini-Mental State Examination (MMSE) (*p* 0.4) or deficit of eye-movement, one of the most typically impaired motor functions in PSP patients (Litvan et al., 1996; Steele et al., 1964) (*p* > 0.2) (Fig. 4B, red). The degree of tau deposition in the GP, in contrast, significantly explained the deficit in eye-movement (*p* < 0.03, FDR-corrected), but did not explain MMSE or any of the above frontal cognitive scores (*p* > 0.2) (Fig. 4B, blue). Similar pattern was also observed for tau deposition in the midbrain.

**Fig. 4.**
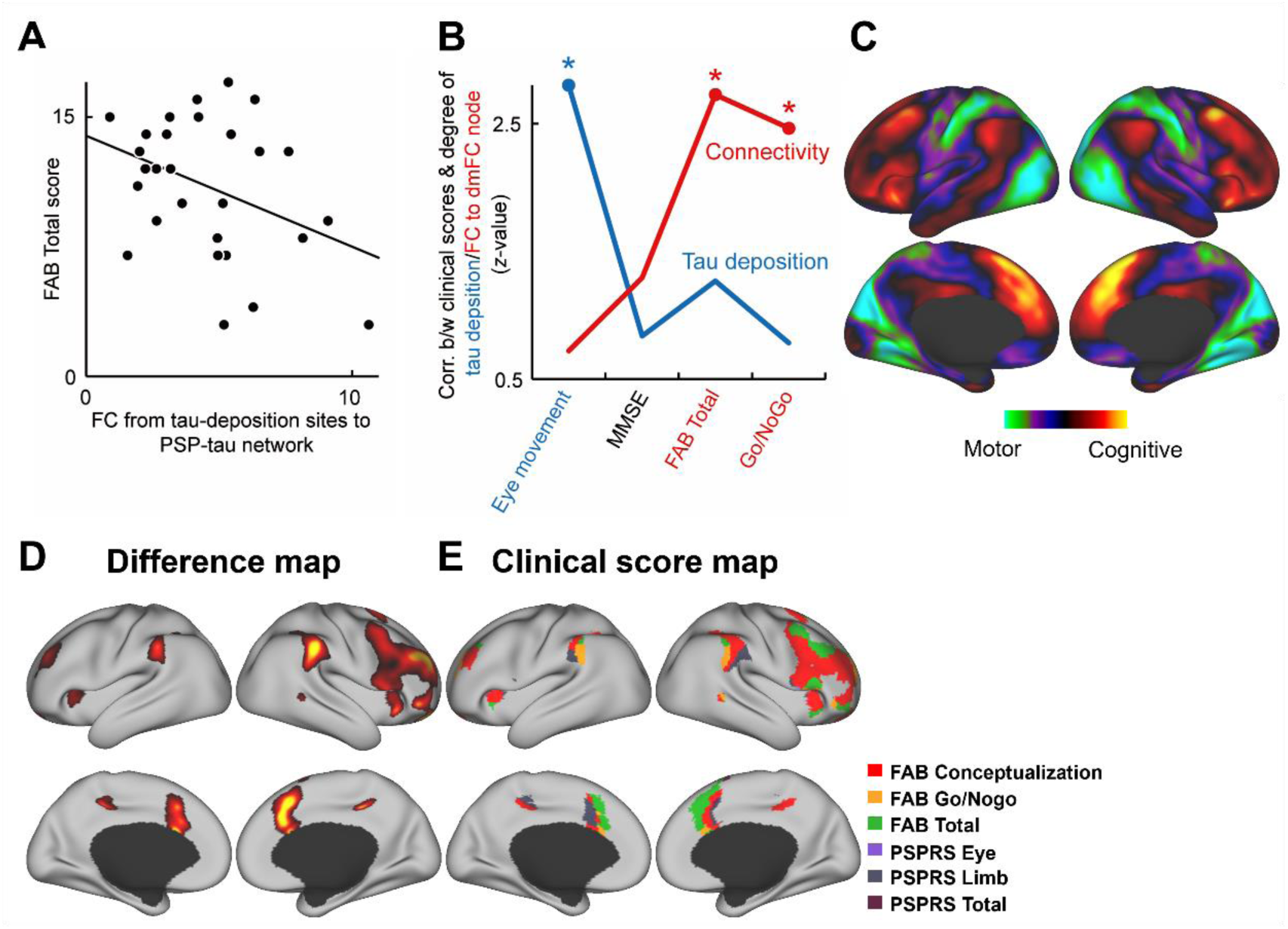
FC strength from tau-deposition sites to the PSP-tau network explained across-patient variability in frontal cognitive decline. **(A)** Scatter plot showing that the FC value from tau-deposition sites to the PSP-tau network exhibited significant negative correlation with the FAB total score across the patients (*p* < 0.02). **(B)** Correlations between motor/frontal-cognitive clinical scores and the degree of tau deposit in GP (blue) or FC value from tau-deposition sites to dmFC node of the PSP-tau network (red). *; *p* < 0.05, FDR-corrected for multiple comparisons. **(C)** Brain maps showing the regions whose FC values from tau-deposition sites primarily explained motor (PSPRS limb) or frontal-cognitive (total FAB score) dysfunction. **(D and E)** The clinical scores with the strongest correlations were assigned to the brain regions in the PSP-tau network (D, same as Fig. 3C) using a winner-take-all method (E).

To map the regions whose FC from the tau-deposition sites predominantly explain the frontal cognitive impairments compared to the motor symptoms among the whole brain, we computed a map that contrasted the FC value–frontal cognitive score correlations with the FC value–motor score correlations (Fig. 4C). After computing correlation maps for each of the FAB/PSPRS scores and their sub-scores (Fig. S7), the scores showing the strongest association with FC values (total FAB score and the PSPRS sub-score of limb movement deficit) were adopted to represent frontal cognitive and motor symptoms. The brain areas whose FC from the tau-deposition sites predominantly explained the frontal cognitive deficits were the adlPFC, dACC, premotor cortex, IFC, AI, and PPC, all of which were consistent with the PSP-tau network (Figs. 4C and 4D) despite that these analyses were operationally independent of each other. Indeed, the above contrast map significantly correlated with the PSP-tau network (*r* = 0.78, *p* < 0.001). Only the correlation maps of frontal cognitive scores (i.e., FAB total, conceptualization, and Go/NoGo task) highly overlapped with the PSP-tau network. To visualize the most relevant clinical score for each area in the PSP-tau network, we created a clinical score map by assigning the score showing the highest positive or negative correlation with the FC values (Fig. 4E). Across the PSP-tau network nodes, FC from tau deposition sites in individual patients was more strongly associated with frontal cognitive impairments than with motor deficits.

Together, these results suggest that at the single-patient level, while the motor symptoms result directly from the local effect of subcortical tau deposits, the frontal cognitive symptoms can be best explained by remote effects of tau deposits exerted on the PSP-tau network.

## DISCUSSION

Here, we conducted tau- and atrophy-network mapping for PSP patients based on multimodal imaging with a tau PET using our recently developed high-contrast tracer florzolotau and structural MRI. While tau deposits and atrophy were primarily detected in subcortical regions, the derived PSP-tau network was predominantly distributed across AMN and FPN. This network was derived specifically from tau-deposition, but not from atrophy. Critically, while the across-patient variance in motor symptoms was significantly explained directly by local effects of subcortical tau deposits, the frontal cognitive symptoms were significantly explained only by remote effects of tau deposits exerted on the cortical PSP-tau network. These results suggest that the functional impacts of tau deposits in heterogeneous regions across PSP patients remotely converge onto the cortical PSP-tau network, leading to their characteristic frontal cognitive symptoms.

Network mapping with normative rs-fMRI data from a large population of healthy individuals has been extensively conducted for brain lesions especially from stroke patients (Siddiqi et al., 2022), and recently applied to atrophy (Tetreault et al., 2020) and tau deposits (Luan et al., 2024) in AD patients. The present study effectively extended this approach to a non-AD tauopathy, with a demonstration of critical distinctions between the efficacy of tau deposits and atrophy in the network mapping, and between the influence of local tau deposits and their remote network effects on specific symptoms for the first time. In particular, the severity of frontal cognitive symptoms, but not motor symptoms, in individual patients could be specifically explained by the FC strength from the tau-deposition sites to the derived PSP-tau network, a proxy for the degree to which tau deposits exert functional impact on these remote regions in each patient. In contrast, local amount of tau in the subcortical regions where both the *t*-value and the overlap across patients were prominent could only predict motor deficits. These results demonstrate the significance of tau-network mapping for elucidating brain-symptom relationships in patients with non-AD tauopathy. The current results indicating remote effects of tau deposits on particular symptoms is consistent with the notion that cortical dysfunctions in PSP patients do not result directly from local tau deposits, but rather stem from network disruption that are remotely triggered by subcortical tau accumulation (Roemer et al., 2024). We also confirmed that even PSP-tau network calculated by seeding only from subcortical tau-deposition sites primarily composed of the same canonical networks as those for the original PSP-tau network. Although the current use of tau-network mapping itself did not specifically focus on cognitive deficits, the resultant strength of the FC from tau-deposition sites to the derived PSP-tau network specifically explained frontal cognitive aspects of the symptoms, rather than motor dysfunctions or general cognitive disabilities. This is likely because the derived PSP-tau network included frontal cortical nodes and largely overlapped with the AMN and FPN as discussed in a later section.

Tau deposition sites were not entirely heterogeneous across the PSP patients, but were partially overlapped, especially in the GP and midbrain. Indeed, these regions have been shown to exhibit significant positive FC and direct/indirect anatomical connectivity with some nodes in the PSP-tau network (Gardner et al., 2013; Middleton and Strick, 1994). This might raise a concern that the derived PSP-tau network merely reflected the FC map seeded from a specific primary region of tau deposit. However, correlations with clinical scores showed substantially different patterns between the amount of tau deposited in each of these regions and the strength of the FC from the tau-deposition sites to the PSP-tau network (Fig. 4). This suggests that the derived PSP-tau network was not merely a simple reflection of the FC pattern from a specific single brain region with highly overlapped tau depositions across patients, but instead presumably reflected the FCs from multiple nodes where tau deposits were distributed heterogeneously across the patients.

While tau-network mapping revealed a prominent convergent cortical network, atrophy-derived similar approach for the same patients did not, even across a range of statistical thresholds for defining seed regions. Note, however, that atrophy-network mapping with a liberal threshold for defining seed regions significantly detected AI region, one of PSP-tau network nodes, as a convergent cortical area commonly connected from atrophy sites across the patients (fig. S3G), suggesting the effectiveness of atrophy-network mapping and a commonality between the resultant network maps derived from tau deposition and atrophy. Overall, the results suggest a potential advantage of using tau deposits detected by florzolotau PET instead of atrophy captured by structural MRI to map a common core pathological network across PSP patients and to clarify the brain-symptom relationships. Tau deposition itself should be toxic to neurons directly and indirectly, and thus would lead to neuronal dysfunctions even without atrophy. The current results suggest the functional impacts of tau deposition itself via remote common network in the PSP. Note that atrophy-network mapping itself has been successfully conducted in previous studies including those for more large cohorts of patients with AD (Tetreault et al., 2020) or schizophrenia (Makhlouf et al., 2024). Therefore, more elaborated analysis of structural MRI data including normative modeling (Segal et al., 2023) with a larger cohort might improve the resultant atrophy-network map for the PSP in future research.

Previous studies have shown significant changes of functional and anatomical connectivity in patients with PSP compared to healthy controls. In particular, cortico-subcortical changes in FC, including those between the midbrain/thalamus and cortical areas, have been reported (Gardner et al., 2013; Piattella et al., 2015). Cortical targets of such FC changes partially overlap with the currently identified PSP-tau network. However, relationships between such cortico-subcortical FC changes and canonical functional networks or frontal cognitive symptoms have not been examined, and the mechanistic difference between motor and frontal cognitive symptoms have not been suggested. Furthermore, significant hypoperfusion and hypometabolism have been detected in the dACC, midcingulate cortex, and dlPFC of patients with PSP (Chiu et al., 2012; Salmon et al., 1997; Varrone et al., 2007), which located in spatial proximity to a subset of the PSP-tau network nodes. Therefore, these physiological changes in PSP patients might at least be partly explained by the tau deposition-derived extended network found in the present study.

Among the canonical networks, the PSP-tau network was most largely overlapped with AMN (Dosenbach et al., 2025), which is associated with planning, execution, monitoring, stopping of action, and attentional allocation. These functions of the network are well consistent with major symptoms of the PSP. Note that, however, the PSP-tau network was also overlapped with a part of FPN, with which AMN jointly supports adaptive and goal-directed behavior (Marek and Dosenbach, 2018). This indicates that the PSP-tau network is not merely a reflection of a single canonical network, but rather a composite of a few distinct networks, thereby exhibiting unique characteristics and association with clinical symptomatology. Critically, FC strength in individual patients between tau-deposition sites and the dmFC node in the PSP-tau network significantly explained the performance of a Go/NoGo task (Fig. 4), which requires the stopping of planned actions. The deficit in response inhibition is one of primary symptoms of the PSP (Migliaccio et al., 2020; Zhang et al., 2016), and is likely reflected in the declined performance of a Go/NoGo task. Furthermore, not only the dmFC node, but a majority of nodes in the PSP-tau network, are recruited by tasks requiring response inhibition (Aron and Poldrack, 2006), and the across-patient correlation map between the Go/NoGo task performance and the FC strength from tau-deposition sites indeed highly overlapped with the whole PSP-tau network. These results suggest that the PSP-tau network is composed of a unique set of AMN/FPN nodes, characterizing the frontal cognitive symptoms of PSP, especially impaired response inhibition.

The present study did not examine whether subcortical tau depositions cause actual dysfunction of the cortical PSP-tau network or whether dysfunction of the PSP-tau network induces frontal cognitive symptoms. Recently, manipulation of neural activity has become possible in non-human primates using chemogenetic techniques such as Designer Receptors Exclusively Activated by Designer Drugs (DREADDs) and Pharmacologically Selective Actuator Module / Pharmacologically Selective Effector Molecule (PSAM/PSEM) systems (Hori et al., 2023; Nagai et al., 2020), and remote effects of neuronal silencing have been visualized at functionally connected regions with behavioral relevance (Hirabayashi et al., 2024; Hirabayashi et al., 2021). Therefore, causal validation and mechanistic understanding of the results obtained with tau-network mapping via reverse translational interventions in non-human primates is an intriguing direction for future research.

Other major limitations of the present study are as follows. First, although within-cohort reproducibility of the PSP-tau network was confirmed in a control analysis, its validation across independent cohorts would be required for more rigorous testing of reproducibility. Another limitation is the cross-sectional design of the research. Longitudinal study of the tau-PET and tracking changes in the PSP-tau network in association with the assessment of symptom progression would lead to a more causal understanding of the relationships between the PSP-tau-network and frontal cognitive symptoms. Another possibility for causal analysis is to test whether neuromodulation that targets a node within the PSP-tau network can reduce frontal cognitive symptoms. If effective, it would represent a novel treatment option against PSP that relies on biological evidence.

The present study demonstrated that the tau-network mapping was able to identify a common cortical network to which tau deposition sites across PSP patients are functionally connected. Tau-PET images obtained from PSP patients can thus be applied to identify the relationships linking the predominantly subcortical tau deposits with the remote dysfunction of the connected cortical networks and the resultant frontal cognitive symptoms in individual patients using normative connectome data. The current approach provides a potential target for symptom-specific neuromodulatory treatment for patients with PSP, and might inform the future risk of frontal cognitive decline for patients in the early phase of the disease, based on still subthreshold amounts of tau deposits at brain regions connected to the PSP-tau network. Furthermore, this approach would be also widely appliable to other tauopathies including traumatic brain injury (Takahata et al., 2019) and late-onset psychiatric disorders (Kurose et al., 2025), for which tau deposits can be reliably detected with florzolotau, as well as for other proteinopathies such as α-synucleinopathies (Endo et al., 2024), or other PET ligands measuring synaptic density or inflammation, to understand the mediating networks between the PET-detectable local pathological changes and the resultant symptoms in a range of brain diseases.

## MATERIALS AND METHODS

### Participants

Forty-six patients with PSP-Richardson syndrome (PSP-RS) and 50 healthy controls (HCs) aged older than 40 years, without histories of neurological disorders were recruited from our affiliated hospitals and the volunteer association of our institute, respectively. Patient details were described in our previous paper (Endo et al., 2022), but PSP participants with PSP-tau scores (Endo et al., 2022) less than 0.3 and HCs with PSP-tau scores more than 0.1 were excluded from the later analyses. The final numbers of participants included were 37 and 48 for patients and HCs, respectively. The participants underwent several neurological examinations, including the PSPRS, FAB, and MMSE.

### PET and MRI data acquisition and preprocessing

Florzolotau PET images were acquired to detect tau deposits using a Biograph mCT flow system (Siemens, Erlangen, Germany) with 2 mm isotropic voxels. T1-weighted images (T1w) were acquired using a 3T MRI scanner (MAGNETOM Verio, Siemens, Erlangen, Germany) for registration of PET images to the standard MRI space and evaluating atrophy (repetition time [TR] = 2300 ms; echo time [TE] = 1.95 ms; inversion time [TI] = 900 ms; flip angle = 9°; field of view [FOV] = 250 mm; matrix size = 512 × 512 × 176; slice thickness = 1 mm). Detailed preprocessing was described in our previous paper (Endo et al., 2022). Briefly, standardized uptake value ratio (SUVR) images were generated from averaged PET images for each participant with motion correction at the intervals of 90–110 min after intravenous injection of ^18^F-florzolotau (186 ± 8.4 [MBq]). Individual SUVR images were spatially normalized to the standard Montreal Neurological Institute (MNI) space (East Asian brain T1w image from the International Consortium for Brain Mapping) using the Diffeomorphic Anatomical Registration Through Exponentiated Lie Algebra (DARTEL) algorithm from Statistical Parametric Mapping (SPM12, Wellcome Department of Cognitive Neurology), running on MATLAB (MathWorks).

### Tau-network mapping

Tau-network mapping is an approach based on lesion-network mapping that identifies brain regions functionally connected to lesion locations. It does not use rs-fMRI scans of the participants themselves, but instead uses a large-scale open database of rs-fMRI acquired from healthy individuals. In this study, FC maps were computed using rs-fMRI data obtained from 100 healthy participants (Boes et al., 2015) from the Brain Genomics Superstruct Project (GSP) (Yeo et al., 2011) with the locations of tau deposits in individual patients as seed regions. Fig. S2 shows the analytical pipeline for tau-network mapping. In the first step, the regions of tau deposits for each patient were identified by comparisons with tau PET images of healthy controls. The *t*-score maps for individual patients were thresholded at *p* < 0.001 (two-sided, uncorrected), and were binarized to create the volumes of interest (VOIs) for calculating FC maps. Next, the mean timeseries from the tau-deposit locations (i.e., VOIs) of a given patient for rs-fMRI data of each participant were compared with the timeseries from every voxel to calculate the correlation map. The maps for all 100 rs-fMRI datasets were then combined to calculate the *t*-score map for each patient. As a control condition of tau-network mapping, we also calculated *t*-score maps using pseudo-randomized VOIs. Briefly, the seeds with the same number of voxels as those of the original tau depositions were randomly selected across the whole brain. The *t*-score maps were then calculated using these pseudo randomized VOIs in the same manner as the original *t*-score maps for tau deposits. Finally, random control-subtracted tau-derived FC map was obtained by contrasting between the FC maps derived from tau deposits and the randomized seeds. FC map calculation, statistical tests, and surface mapping were performed using FMRIB Software Library’s (FSL’s) FEAT tool (Smith et al., 2004), Analysis of Functional NeuroImages (AFNI) software (Cox, 1996), and Connectome Workbentch (Marcus et al., 2011), respectively.

### Atrophy-network mapping

Data preprocessing for VBM was performed using SPM12 on MATLAB and PMOD 4.2 / 4.3 (PMOD Technologies LLC, Switzerland). Gray and white matter segmentation images were extracted from T1w images and combined to obtain the parenchymal image using SPM12. The DARTEL algorithm was then employed to spatially normalize each brain parenchymal image (gray and white matter) by preserving amounts in MNI space. The regions of atrophy for each patient were identified by comparisons with parenchymal images of healthy controls. The *t*-score maps for each patient were thresholded at *p* < 0.001 (two-sided, uncorrected), and were binarized to create the VOIs for calculating FC maps. Later analyses were the same as those for the tau-network mapping described above.

### Assignments of canonical networks to the derived PSP-tau network

Based on the atlas in Ji et al.(Ji et al., 2019), one of twelve canonical networks including the orbito-affective, ventral multimodal, posterior multimodal, default mode, auditory, frontoparietal, language, dorsal attention, cingulo-opercular [action-mode (Dosenbach et al., 2025)], somatomotor, and primary and secondary visual networks were assigned to each brain area in the tau-network map before comparison with atrophy-network map and the PSP-tau network that was determined as a significant (*p* < 0.05, FWE-corrected) regions in a statistical comparison (paired *t*-test) between the FC maps derived from tau- and atrophy-network mapping among the patients. Furthermore, we also assigned the canonical networks to each brain area in the tau-network map, calculated based solely on subcortical tau deposits across all patients. The proportion of each network among all areas was then calculated and statistically evaluated using a Chi-square test. To identify the SCAN within the SMN, we used three coordinates of inter effectors in the superior, middle, and inferior parts of the primary motor areas reported by Gordon et al (Gordon et al., 2023).

### Validation analyses of tau/atrophy-network mapping

To examine the consistency/stability of the tau/atrophy-network mapping results among different criteria for the seed definition, both more liberal (*p* < 0.005 uncorrected) and more stringent (*p* < 0.05, FWE) thresholds compared with the original one (*p* < 0.001) were also tested to identify the locations of tau deposition or atrophy in the patients for subsequent network mapping. To confirm the reproducibility of tau/atrophy network mapping in the current cohort of patients, tau/atrophy-network maps calculated for individual patients (N = 37) were randomly divided into two groups 100 times, and the tau/atrophy network mapping was performed for these two groups separately. Then, the resultant FC values (from the tau-deposition sites) in the Johns Hopkins University (JHU) whole brain VOIs were compared between the two groups. We assessed statistical significance by testing whether the mean correlation coefficients between the two groups exceeded the 95th percentile of the correlation coefficients obtained for pairs of randomly shuffled ROIs (fig. S4, D and H). To confirm the consistency of tau-network mapping results among different rs-fMRI databases, we also performed tau-network mapping based on the Human Connectome Project (HCP) (Van Essen et al., 2013) data. For each patient, the FC map seeded from the tau-deposition sites was calculated with the HCP young adult dataset (N = 98) in the same manner as the analysis using the GSP dataset described above. The voxels with top 10% FC values in the cortex were then extracted from the resultant tau-network maps based on GSP/HCP data for comparison between the two rs-fMRI datasets. FC values in the JHU whole brain VOIs were also calculated to quantify the threshold-free correlation between the maps obtained with GSP/HCP data.

### Across-patient correlation between clinical scores and FC values from tau-deposition sites to the PSP-tau network

To complement the single regression analysis showing that the FAB total score could be significantly explained by the FC strength from tau deposition sites to the PSP-tau network (Fig. 4A), multiple regression analysis was further conducted using an R package, in which additional control explanatory variables were the degree of tau deposition in the GP, where both the maximum overlap and the highest *t*-value of tau deposition were observed, and in the PSP-tau network itself. To more closely examine the relationships between the severity of frontal cognitive/motor symptoms and the remote/local effect of tau deposition across patients (Fig. 4B), we then extracted the FC strength from tau-deposition sites to the dmFC node of the PSP-tau network, a representative frontal node in the network and the amounts of tau deposition in the GP. Pearson’s correlation coefficients were then calculated between these values and clinical scores (PSPRS and FAB) including their sub-scores. MMSE served as a control score reflecting general cognitive ability. Statistical significance was assessed using false discovery rate (FDR) correction for multiple comparisons. In order to visualize the regions whose FC from tau-deposition sites predominantly explained the frontal cognitive impairments against motor-related dysfunctions at the whole-brain level (Fig. 4C), we contrasted the whole-brain correlation map between the FC and the frontal cognitive symptoms against that between the FC and motor deficits. Based on the maximum values of correlation coefficient between the FC strength and clinical scores, total score of the FAB and limb deficit score of the PSPRS were adopted as indices representing frontal cognitive and motor impairments, respectively. Finally, to visualize the clinical score that most strongly correlated with the FC value from tau deposition sites to each brain region within the PSP-tau network (Fig. 4E), the whole-brain correlation maps between the FC value from tau deposition sites and individual clinical scores were calculated, and the clinical scores with the strongest correlation were assigned to each brain region of the PSP-tau network. The above whole brain analyses were performed using SPM12, Connectome Workbench, and FSL software.

## Acknowledgments

We thank all patients and their caregivers, as well as volunteers, for their participation in this study; clinical research coordinators; PET and MRI operators; radiochemists; and research ethics advisers at Quantum Science and Technology (QST) for their assistance with the current projects. We acknowledge the support of Matsuoka, K. Hirata, M. Oya, H. Matsumoto, M. Ichihashi, Y. Komatsu, Y. Yamamoto, M. Kubota, K. S. Kitamura, Y. Takado, K. Kawamura, and M.R. Zhang at QST. We thank APRINOIA Therapeutics for kindly sharing the precursor of florzolotau (18F). We acknowledge the support of S. Hirano and Y. Nakano at the Department of Neurology, Chiba University on patient recruitment; T. Hatano, T. Tsunemi, N. Nishikawa, K. Nishioka (currently working at The Juntendo Tokyo Koto Geriatric Medical Center), Y. Yamashita, Y. Motoi, and S. Saiki (currently working at the University of Tsukuba) at the Department of Neurology, Juntendo University School of Medicine; I. Aiba at the Department of Neurology, National Hospital Organization Higashinagoya National Hospital; T. Yuasa at the Department of Neurology, Kamagaya General Hospital; H. Imai at the Tokyo Rinkai Hospital; Y. Nishida and Y. Yagi at the Department of Neurology, Tokyo Medical and Dental University; S. Furukawa at the Narita Red Cross Hospital; M. Seki at the Department of Neurology, Keio University School of Medicine; and T. Takeda and I. Isose at the Department of Neurology, Chiba-East Hospital. Data were provided in part by the Human Connectome Project, WU-Minn Consortium (Principal Investigators: David Van Essen and Kamil Ugurbil; 1U54MH091657) funded by the 16 NIH Institutes and Centers that support the NIH Blueprint for Neuroscience Research; and by the McDonnell Center for Systems Neuroscience at Washington University. We also thank Adam Phillips, PhD, from Edanz (https://jp.edanz.com/ac) for editing a draft of this manuscript.

## Funding

AMED Grant JP24wm0625307 (to TH)

AMED Grant JP19dm0207072 (to MH)

AMED Grant JP24wm0625001 (to MH)

JST Grant JPMJMS2024 (to MH)

MEXT/JSPS KAKENHI JP24H00734 (to TH)

MEXT/JSPS KAKENHI JP25H01767 (to YH)

MEXT/JSPS KAKENHI JP23K11796 (to YH)

Biogen Idec Inc.

APRINOIA Therapeutics.

## Author contributions

Conceptualization: TH

Methodology: YH, TH, HE

Investigation: YH, TH, HE

Visualization: YH, TH

Funding acquisition: TH, YH, HE, MH, TM

Project administration: TH

Supervision: TH, EH, TM, MH

Writing – original draft: YH, TH

Writing – review & editing: All authors

## Data and materials availability

The data supporting our study findings are available from the corresponding author upon reasonable request.

