## Supplemental Information for "Remote Cortical Network for Frontal Cognitive Symptoms Derived from Tau Accumulation in Progressive Supranuclear Palsy"

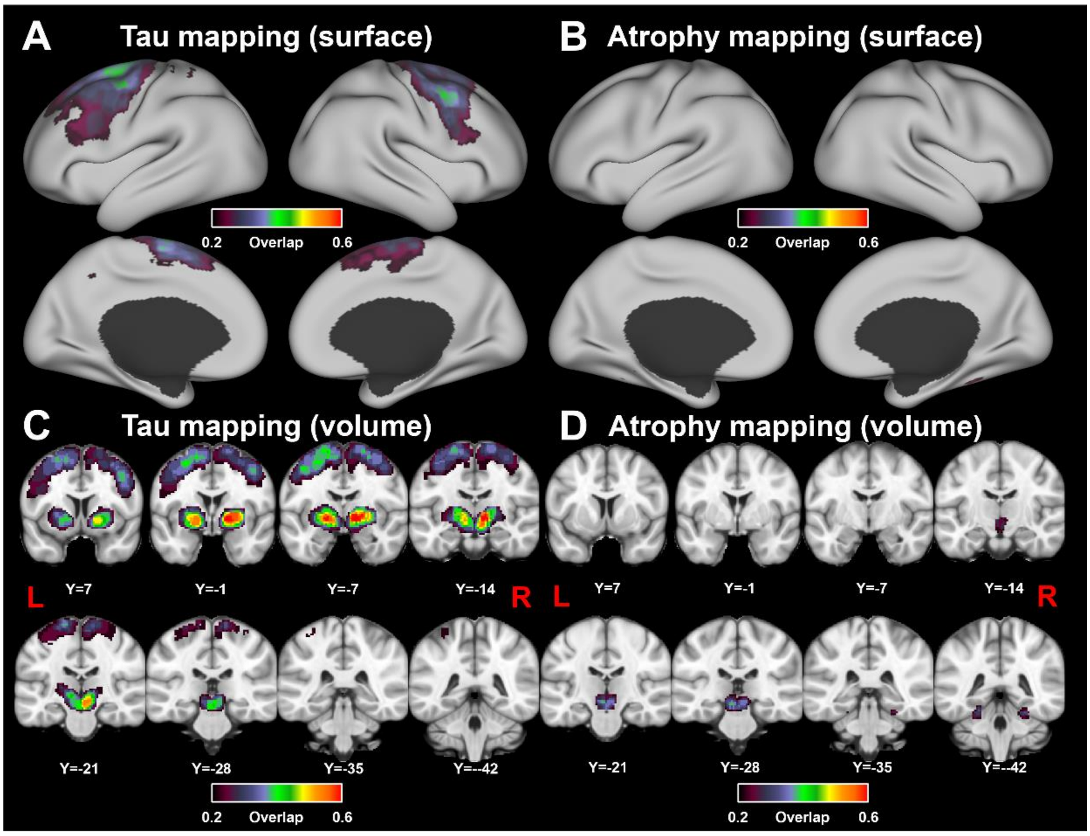


**fig. S1 Overlap of tau deposit and brain atrophy locations among patients with PSP**. Overlap maps of tau deposit (A, C) and atrophy (B, D) are presented on the surface (A, B) and in volume space (C, D). Significant tau accumulation was observed in the GP (thresholded at $p < 0.001$, overlapped maximally for 62% of patients), thalamus (62%), midbrain (57%), and some regions in the motor-related cortex (25%), while brain atrophy was significantly observed in the midbrain (35%) and cerebellum (32%).

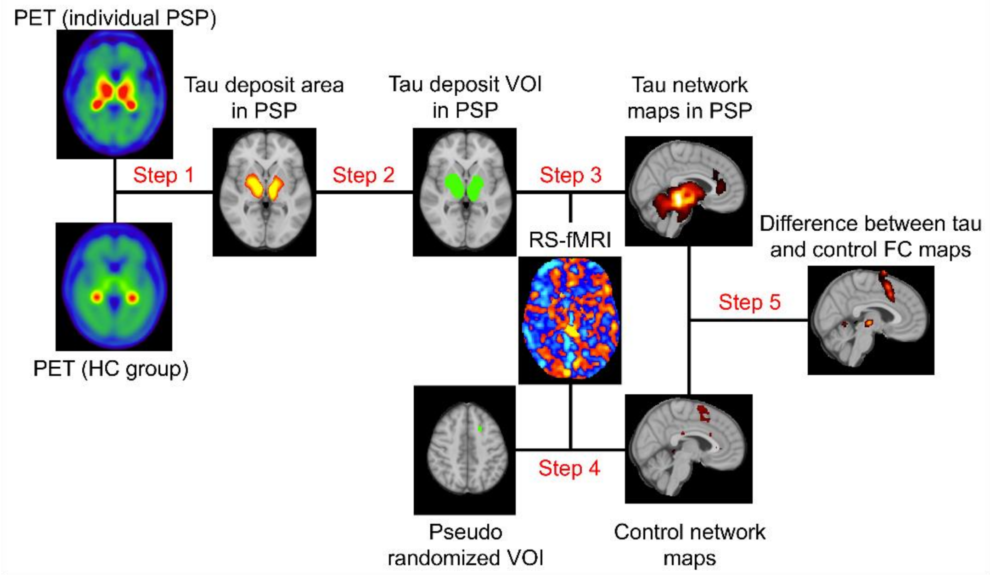


**fig. S2 Pipeline for tau-network mapping**. Tau deposition sites for each patient were identified by comparisons with tau-PET images from healthy controls (Step 1). The *t*-score maps of tau deposition for individual patients were then thresholded at $p < 0.001$ (two-sided, uncorrected), and binarized to create the seed VOIs (Step 2). Next, the mean timeseries in the VOI from each rs-fMRI data ($N = 100$) were compared with timeseries for each voxel to calculate the correlation maps (tau-network maps). The resultant maps for all 100 rs-fMRI datasets were then combined to calculate the *t*-score map for each patient (Step 3). As a control condition, *t*-score maps were calculated using pseudo-randomized seed voxels for individual patients (Step 4) Finally, tau-derived FC map was obtained by calculating a differential *t*-score map between tau deposit-derived and control FC maps (Step 5).

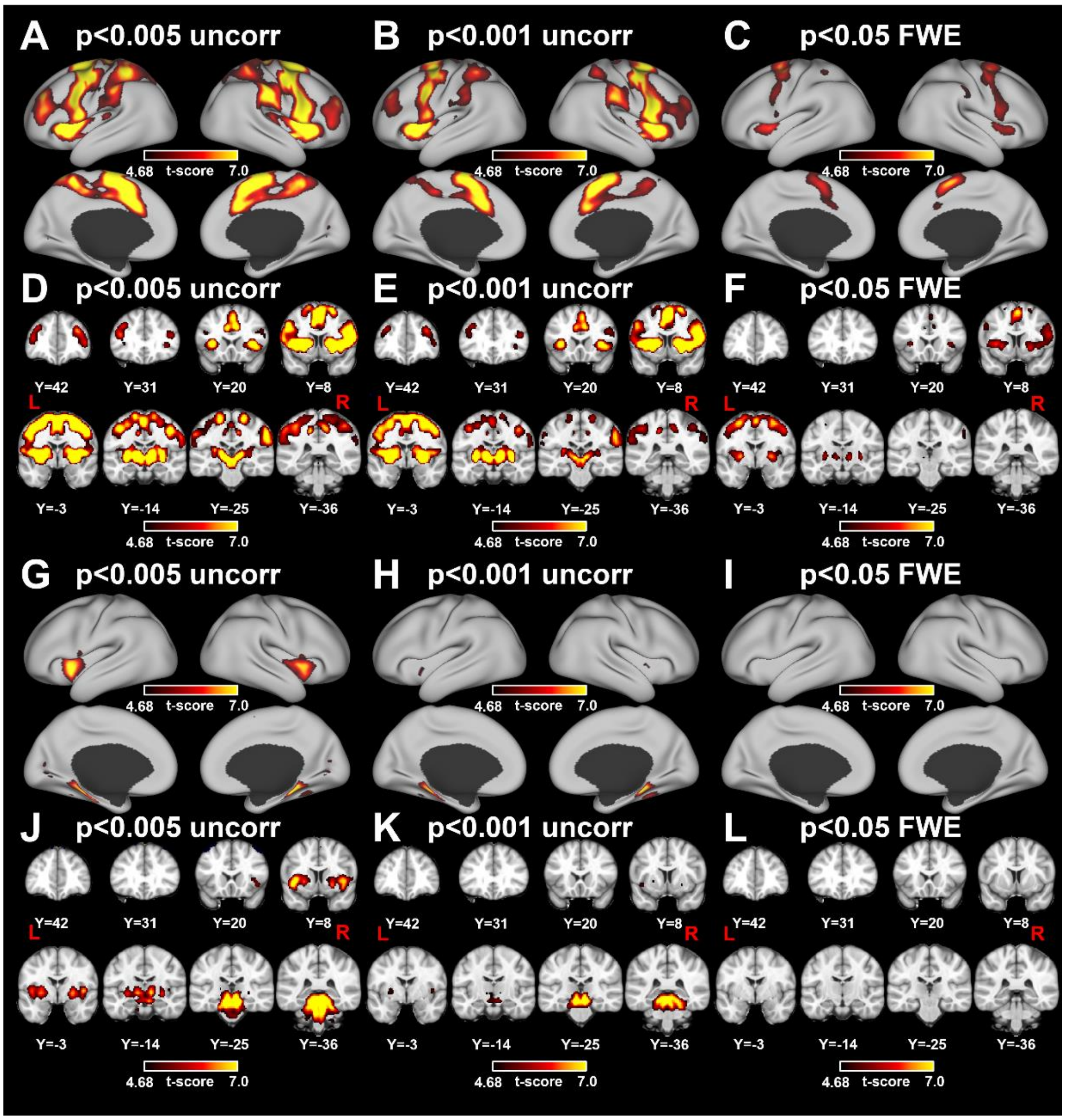


**fig. S3 Tau/atrophy-network maps with different thresholds for seed definition.** Tau (A-F) and atrophy (G-L) network maps were obtained using seed regions derived from tau deposits and atrophy with three different thresholds ($p < 0.005$ uncorrected, $p < 0.001$ uncorrected, and $p < 0.05$ FWE-corrected).

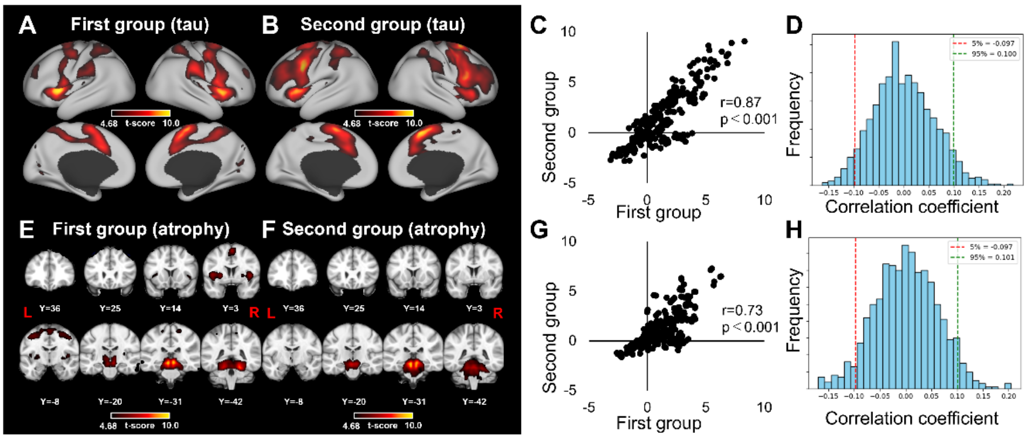


**fig. S4 Reproducibility of tau/atrophy-network mapping. (A-D)** Comparisons of tau-network mapping results for split-half subgroups. Correlation of FC values for the same region was plotted in (C). The histogram of correlation coefficients obtained from randomly shuffled ROIs showed that the 5th and 95th percentile values were -0.097 and 0.10, respectively (D). **(E-H)** Same as (A-D), but for atrophy. Volume maps were adopted instead of surface maps for atrophy due to subcortical predominance. The VOI values in the tau/atrophy-network maps across the whole brain were highly consistent between the first and second groups of patients ($r$ = 0.87 and 0.73, $p < 0.001$ for tau- and atrophy-network map, respectively). When all patients were randomly divided into two groups 100 times, the mean correlation coefficient ($r$ = 0.92 and 0.88, $p < 0.001$ for tau- and atrophy-network map, respectively) far exceeded the 95th percentile of the correlation coefficients obtained from randomly shuffled ROIs.

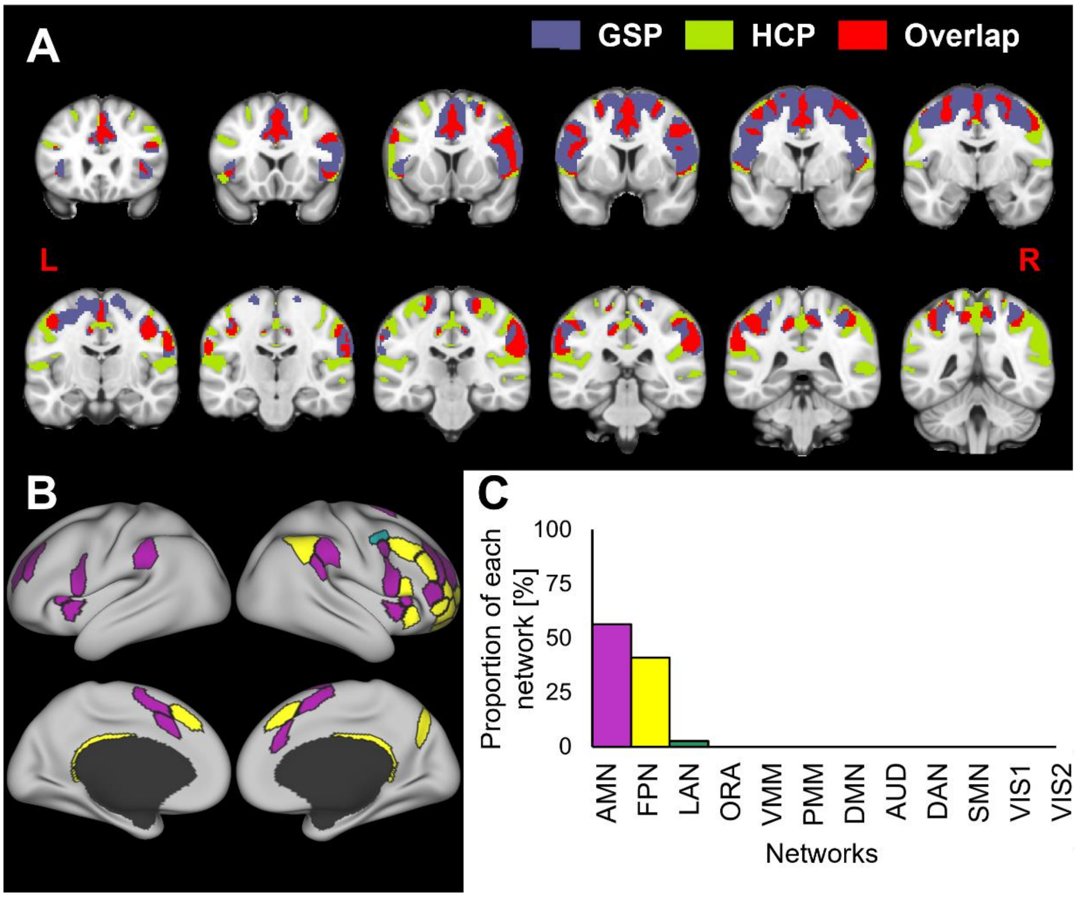


**fig. S5 Consistency of tau-network mapping results across two different connectomes**. **(A)** Tau-network maps were computed based on GSP and HCP databases. Voxels showing top 10% of the FC values in the cortex are shown in blue and green for GSP and HCP, respectively. Overlap between the tau-network maps based on the two databases were depicted in red, which included the LPFC, dACC, PPC, and AI. **(B and C)** Similar to Fig. 3H, one of twelve canonical networks was assigned to each region in the tau-network maps computed using HCP database (B), and the proportion of each network area to the total area was calculated (C).

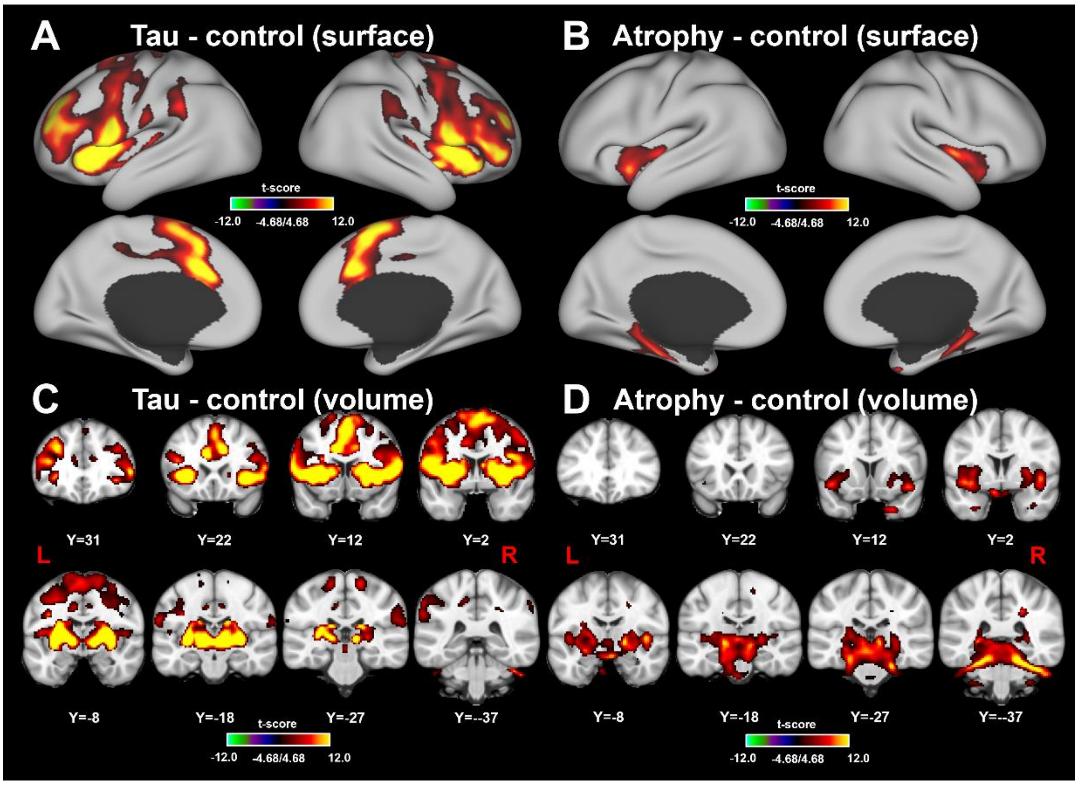


**fig. S6 Comparison of tau/atrophy-network maps with control FC maps calculated with spatially pseudo-randomized seed regions. (A and C)** Control-subtracted tau-network map presented in the surface (A) and volume (C) spaces, respectively. **(B and D)** Same as (A and C), but for atrophy-network map.

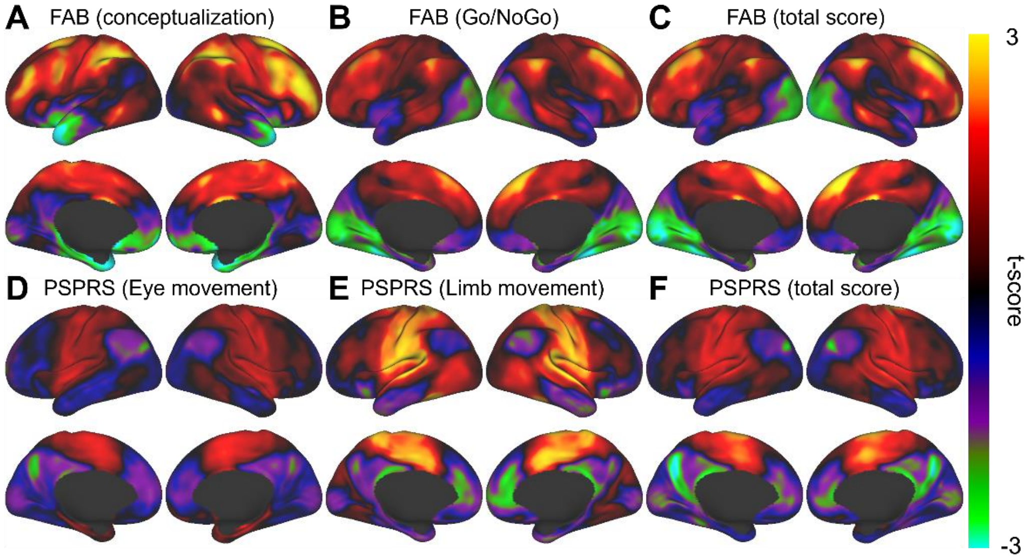


**fig. S7 Correlation maps between the FC value from tau-deposition sites and clinical scores related to frontal cognitive (A-C) and motor (D-F) deficits.**
